# Photobody formation spatially segregates two opposing phytochrome B signaling actions to titrate plant environmental responses

**DOI:** 10.1101/2023.11.12.566724

**Authors:** Ruth Jean Ae Kim, De Fan, Jiangman He, Keunhwa Kim, Juan Du, Meng Chen

## Abstract

Photoactivation of the plant photoreceptor and thermosensor phytochrome B (PHYB) triggers its condensation into subnuclear photobodies (PBs). However, the function of PBs remains frustratingly elusive. Here, we found that PHYB recruits PHYTOCHROME-INTERACTING FACTOR5 (PIF5) to PBs. Surprisingly, PHYB exerts opposing roles in degrading and stabilizing PIF5. Perturbing PB size by overproducing PHYB provoked a biphasic PIF5 response: while a moderate increase in PHYB enhanced PIF5 degradation, further elevating the PHYB level stabilized PIF5 by retaining more of it in enlarged PBs. These results reveal a PB-mediated light and temperature sensing mechanism, in which PHYB condensation confers the co-occurrence and competition of two antagonistic phase-separated PHYB signaling actions—PIF5 stabilization in PBs and PIF5 degradation in the surrounding nucleoplasm— thereby enabling an environmentally-sensitive counterbalancing mechanism to titrate nucleoplasmic PIF5 and its transcriptional output. This PB-enabled signaling mechanism provides a framework for regulating a plethora of PHYB-interacting signaling molecules in diverse plant environmental responses.

## INTRODUCTION

A universal architectural hallmark of the cell nucleus is various functionally distinct membraneless subnuclear organelles, collectively referred to as nuclear bodies^1–3^. Nuclear bodies act as liquid- or gel-like biomolecular condensates that are commonly thought to be excluded from the surrounding nucleoplasm via concentration-dependent, energetically favorable liquid-liquid phase separation (LLPS)^3–5^. Many nuclear bodies, such as nucleoli, Cajal bodies, nuclear speckles, paraspeckles, histone locus bodies, and promyelocytic leukemia protein (PML) bodies, are well characterized, and they are associated with diverse basic nuclear functions in transcription, splicing, and DNA replication and repair. Accumulating observations have also documented the nonuniform subnuclear distribution of cell signaling molecules to nuclear bodies, in addition to the surrounding nucleoplasm. For instance, the tumor suppressor p53 is recruited to PML bodies^6^. However, the functional significance of the spatial segregation of signaling molecules into two phase-separated compartments—nuclear bodies and the surrounding nucleoplasm—remains elusive. A better understanding of the function of nuclear bodies in cell signaling requires new genetically tractable models directly connecting nuclear body dynamics to cell signaling. Photobodies (PBs) are plant nuclear bodies defined molecularly by the presence of the photoreceptor and thermosensor phytochrome B (PHYB)^7–9^. Environmental light and temperature cues control PHYB activity and thereby directly regulate both the assembly/dissolution of PBs and PHYB signaling outputs^7,10–12^. As such, PBs provide a unique experimental paradigm for interrogating the general principles of nuclear bodies in cell signaling.

PHYs are evolutionarily conserved red (R) and far-red (FR) photoreceptors in plants^13^. The prototypical plant PHY is a homodimer; each monomer contains an N-terminal photosensory module and a C-terminal output module^13,14^. PHYs can be photoconverted between two relatively stable forms: the R-light-absorbing inactive Pr form and the FR-light-absorbing active Pfr form^13,14^. The genome of the reference plant species *Arabidopsis thaliana* (*Arabidopsis*) encodes five PHYs, PHYA-E, among which PHYB plays a prominent role^15^. The photoconversion of PHYB—either photoactivation by R light or photoinhibition by FR light— alters the Pfr/Pr equilibrium of individual PHYB molecules and the steady-state cellular concentration of the active PfrPfr homodimer, thereby allowing plants to monitor changes in light quality, quantity, and periodicity. The amount of active PHYB can also be modulated by ambient temperature through temperature-dependent thermal reversion from Pfr to Pr^16^, making PHYB a prominent thermosensor in addition to a photoreceptor^11,12^. By controlling the amount of active PHYB, environmental light and temperature cues regulate all aspects of plant development and growth, including germination, seedling establishment, shade avoidance, floral induction, senescence, and immunity^17,18^. PHYB signaling is best studied during *Arabidopsis* seedling establishment, where the perception of environmental cues by PHYB in the epidermal cells of the embryonic leaves (cotyledons) controls the biosynthesis of the mobile growth hormone auxin to regulate seedling morphogenesis, including promoting cotyledon expansion locally, as well as inhibiting the elongation of the embryonic stem (hypocotyl) remotely^17,18^.

The PHYB-containing subnuclear organelle was named the “photobody” by Joanne Chory in reference to its dynamic assembly and dissolution in response to R and FR light^19^. The “photobody” name was also adopted to describe the fascinating blue-light-inducible nuclear bodies containing the blue-light photoreceptor cryptochrome 2 (CRY2)^20^. However, there is still little evidence demonstrating that the CRY2-containing subnuclear foci are the same as the PBs. Co-expressing PHYB-CFP and CRY2-RFP in BY-2 protoplasts showed distinct PHYB-CFP and CRY2-RFP subnuclear compartments with only some containing both photoreceptors^21^. The relationship between the CRY2-containing subnuclear foci and the PHYB-containing PBs in *Arabidopsi*s remains to be examined. The regulation of PBs by R light and temperature has been extensively studied in *Arabidopsis* pavement epidermal cells using fluorescent-protein-tagged PHYB (PHYB-FP). The current model posits that PB formation and maintenance are driven by the condensation of the active PfrPfr form of PHYB. This conclusion is supported by several lines of evidence. First, PB formation and dissolution can be induced by the activation and inactivation of PHYB, respectively. Akira Nagatani, Ferenc Nagy, and Eberhard Schäfer first reported that the photoactivation of PHYB-FP triggers its nuclear accumulation and further compartmentalization into discrete subnuclear speckles (PBs)^22,23^. Conversely, during the light-to-dark transition or upon an FR treatment where PHYB reverts to the inactive form, PHYB-FP moves from PBs into the nucleoplasm^24–27^. Second, the steady-state pattern of PBs, particularly PB size, correlates with the amount of active PHYB^28^. Under intense R light (e.g., 10 μmol m^-2^ s^-1^), where each PHYB molecule stays in the Pfr form at least 50% of the time, PHYB-FP assembles into two to ten large PBs of 0.7-2 μm in diameter^10,24,28,29^. In contrast, under dim R light, where PHYB stays as the Pr most of the time, PHYB-FP localizes to tens of small PBs of less than 0.1-0.7 μm in diameter or it is dispersed evenly in the nucleoplasm^10,24,28^. Consistent with these observations, a constitutively active *phyB* mutant YHB, which carries a Y276H mutation in PHYB’s chromophore attachment domain and locks PHYB in an active form, localizes to a few large PBs even in the dark^30^. In contrast, *phyB* mutants that destabilize the Pfr form, including by removing the disordered N-terminal extension^16,31,32^, promote PB dissolution^28,31–33^. Third, PB formation relies on the dimerization of PHYB’s C-terminal module^32,34,35^. A D1040V mutation in this module, which disrupts PHYB dimerization, abolishes PHYB’s ability to form PBs^36^. Fourth, PHYB can undergo light-dependent LLPS into biomolecular condensates in mammalian cells^37–39^. The intramolecular attributes of PHYB required for PB formation *in vivo*, e.g., the C-terminal module and N-terminal extension, are also essential for PHYB LLPS in this heterologous system^37^. These observations support the idea that PB formation *in vivo* may be propelled by the LLPS of active PHYB alone, although PHYB condensation under the physiological condition is likely more complex involving other molecules such as PHOTOPERIODIC CONTROL OF HYPOCOTYL 1 (PCH1)^26,40^.

PHYB controls diverse developmental responses primarily by regulating a family of basic helix-loop-helix transcription factors called PHYTOCHROME-INTERACTING FACTORs (PIFs)^18,41,42^. In general, PIFs are antagonists of PHYB signaling. For instance, the five prominent PIFs—PIF1, PIF3, PIF4, PIF5, and PIF7—collectively promote hypocotyl elongation^18,41,42^. A central mechanism of PHYB signaling is to promote the degradation of PIFs^18,41,42^. PIF1, PIF3, PIF4, and PIF5 are rapidly degraded during the dark-to-light transition in a PHYB-dependent manner^43–47^. PHYB binds directly to the prototypical PIF, PIF3, to promote its phosphorylation by PHOTOREGULATORY PROTEIN KINASEs (PPKs) and subsequent subsequent ubiquitin-proteasome-dependent degradation^43,48–50^. PIF3 can be ubiquitylated by two E3 ubiquitin ligases, the **c**ullin3-**R**ING E3 ubiquitin **l**igases (CRL) with LIGHT-RESPONSE BRIC-A-BRAC/TRAMTRACK/BROADs (LRBs) as the substrate recognition subunits (CRL3^LRB^), and the cullin1 E3 with EIN3-BINDING F BOX PROTEINs (EBFs) as the substrate recognition subunits (CRL1^EBF^)^49,50^. In addition to regulating PIFs via direct interaction, PHYB also promotes PIF degradation indirectly by inhibiting that factors that stabilize PIFs, such as the CONSTITUTIVE PHOTOMORPHOGENIC 1 (COP1) and SUPPRESSOR OF PHYTOCHROME A-105 proteins (SPAs), which constitutes the substrate recognition subunits of the CRL4^COP1/SPA^ E3 ubiquitin ligases^51–53^. Interestingly, although PIF1, PIF3, PIF4, and PIF5 are subject to PHYB-mediated degradation, they exhibit different accumulation patterns. While PIF1 and PIF3 accumulate only in darkness and become undetectable in the light^29,36^, PIF4 and PIF5 can accumulate to significant levels under prolonged light treatment^36,45,47,54–56^. The accumulation of PIF4 and PIF5 in the light plays an essential role in the response to warm temperatures during the daytime^54,57^. However, the mechanism that stabilizes PIF4 and PIF5 in the light remains elusive.

PBs are considered to be associated with PHYB signaling^7,8^. PB formation correlates with PHYB-mediated light responses such as the inhibition of hypocotyl elongation^24,28,29^. A large collection of *phyB* mutants that are defective in PB formation also show attenuated PHYB signaling^28,30–34^. Recent proteomics studies revealed that many PHYB signaling components reside in PBs, these components include COP1, SPAs, PPKs, PCH1, PCH1-LIKE (PCHL), and transcription regulators in light and photoperiodic signaling, including PIF4, TANDEM ZINC-FINGER-PLUS 3 (TZP), EARLY FLOWERING 3 (ELF3), EARLY FLOWERING 4 (ELF4), and LUX ARRHYTHMO (LUX), lending further support to the functional role of PBs in PHYB signaling^26,58,59^. The current data support the model that PBs play an important role in PIF3 degradation. During the dark-to-light transition, FP-tagged PIF3 localizes to PBs before its degradation^43,60^. Conversely, PIF3 reaccumulation during the light-to-dark transition correlates with the disappearance of PBs^24^. Forward genetic screens have identified three PB-deficient mutants, *hmr* (*hemera*)^29,61,62^, *rcb* (*regulator of chloroplast biogenesis*)^63^, and *ncp* (*nuclear control of pep activity*)^64^, in which PHYB-FP fails to form large PBs and PIF3 degradation is blocked in the light. Moreover, overexpressing PHYB’s C-terminal output module alone, which constitutively localizes to PBs, is sufficient to mediate PIF3 degradation even in darkness^36^.

Despite the accumulating evidence associating PBs with PIF3 degradation, the precise function of PBs remains frustratingly elusive. We still cannot unequivocally conclude that PBs are the sites of PIF3 degradation, and it is still unknown whether PBs also regulate the stability of other PIFs. A major challenge in dissecting the function of nuclear bodies, in general, has been the difficulty uncoupling the functional output of nuclear bodies from that of the surrounding nucleoplasm. Because components can diffuse between the nuclear body compartment and the surrounding nucleoplasm, although manipulating those components may disrupt nuclear body assembly and the functional output associated with the components, such a correlation is usually not sufficient to assign the function exclusively to the nuclear-body compartment. The previous approach of using loss-of-function mutants in PHYB or PHYB signaling was likely to disrupt the PHYB signaling outputs of both PBs and the surrounding nucleoplasm and therefore could not uncouple the signaling actions of the two phase-separated compartments. Another hurdle in studying PIF3 degradation was the difficulty monitoring its PB localization: because PIF3 does not accumulate to a detectable level in the light, FP-tagged PIF3 can only be observed briefly during the dark-to-light transition^43,60^.

To circumvent these obstacles to characterizing PBs, we implemented two major strategic changes in the current study. First, we adopted PIF5 as a model because PIF5 accumulates in the light and therefore could potentially allow us to visualize its localization to PBs in light-grown seedlings^36,45,47,54–56^, though the PB localization of PIF5 had not been reported previously. Second, instead of disrupting PBs using loss-of-function mutants in PHYB or PHYB signaling, we perturbed the PB size by increasing PHYB abundance. Here, we show that PHYB recruits PIF5 to PBs and, surprisingly, that PHYB exerts two opposing functions in both degrading and stabilizing PIF5. Our results reveal that PB formation enables the phase-separation and competition of the two antagonistic PHYB signaling actions. PHYB condensation stabilizes PIF5 in PBs to counteract PIF5 degradation in the surrounding nucleoplasm, thereby enabling a counterbalancing mechanism to titrate nuclear PIF5 and its signaling outputs. This novel PB-mediated PHYB signaling mechanism provides a framework for regulating a plethora of PHYB-interacting signaling molecules to generate diverse environmental responses in plants.

## RESULTS

### Increasing PHYB production alters PB size and dynamics

Instead of interrogating the function of PBs in loss-of-function *phyB* or PHYB signaling mutants, we explored an alternative approach to perturb PBs by increasing PHYB abundance. The current hypothesis is that PBs form via the LLPS of PHYB^37^. Based on the theory of the LLPS, LLPS of PHYB occurs when PHYB accumulates above at a critical concentration. The PHYB LLPS model predicts that the overproduction of PHYB is expected to result only in the growth of PBs without changing the concentration of PHYB in either the PB or the surrounding nucleoplasmic compartment^4,65^. Therefore, if the model were correct, increasing the PHYB level should only enhance the signaling output from PBs while leaving the functional output of the nucleoplasmic PHYB unchanged or less affected. To test this hypothesis, we collected *Arabidopsis* lines expressing various amounts of PHYB. We previously reported the *PBC* line, which overexpresses CFP-tagged PHYB (**P**HY**B**-**C**FP) in the *phyB-9* background^32^. *PBC* exhibits a short hypocotyl phenotype, as it overexpresses PHYB-CFP at a level 65-fold that of the endogenous PHYB (Fig. 1a-c)^32^. To create lines expressing intermediate levels of PHYB between *PBC* and Col-0, we generated new transgenic lines in the *phyB-9* background carrying *PHYB* genomic DNA with a *CFP* sequence inserted immediately before the *PHYB* stop codon. We named them *gPBC* (***g****enomic **P**HY**B**-**C**FP*) lines. Two single-insertion *gPBC* lines, *gPBC-25* and *gPBC-29*, were selected for further analysis. The steady-state levels of PHYB-CFP in *gPBC-25* and *gPBC-29* were about 7- and 40-fold that of the endogenous PHYB level in Col-0, respectively (Fig. 1a-c). Both *gPBC* lines rescued the long-hypocotyl phenotype of *phyB-9*, indicating that PHYB-CFP was functional (Fig. 1a,b). Corroborating the notion that the PHYB response correlates with the PHYB level, the hypocotyl length of *gPBC-25* was in between that of Col-0 and *PBC*, whereas the hypocotyl length of *gPBC-29* was similar to that of *PBC* (Fig. 1a,b). With the two new *gPBC* lines, we had a panel of five lines—*phyB-9*, Col-0, *gPBC-25*, *gPBC-29*, and *PBC*—with PHYB levels ranging from zero to 65-fold that of the wild-type level (Fig. 1a-c).

**Fig. 1.**
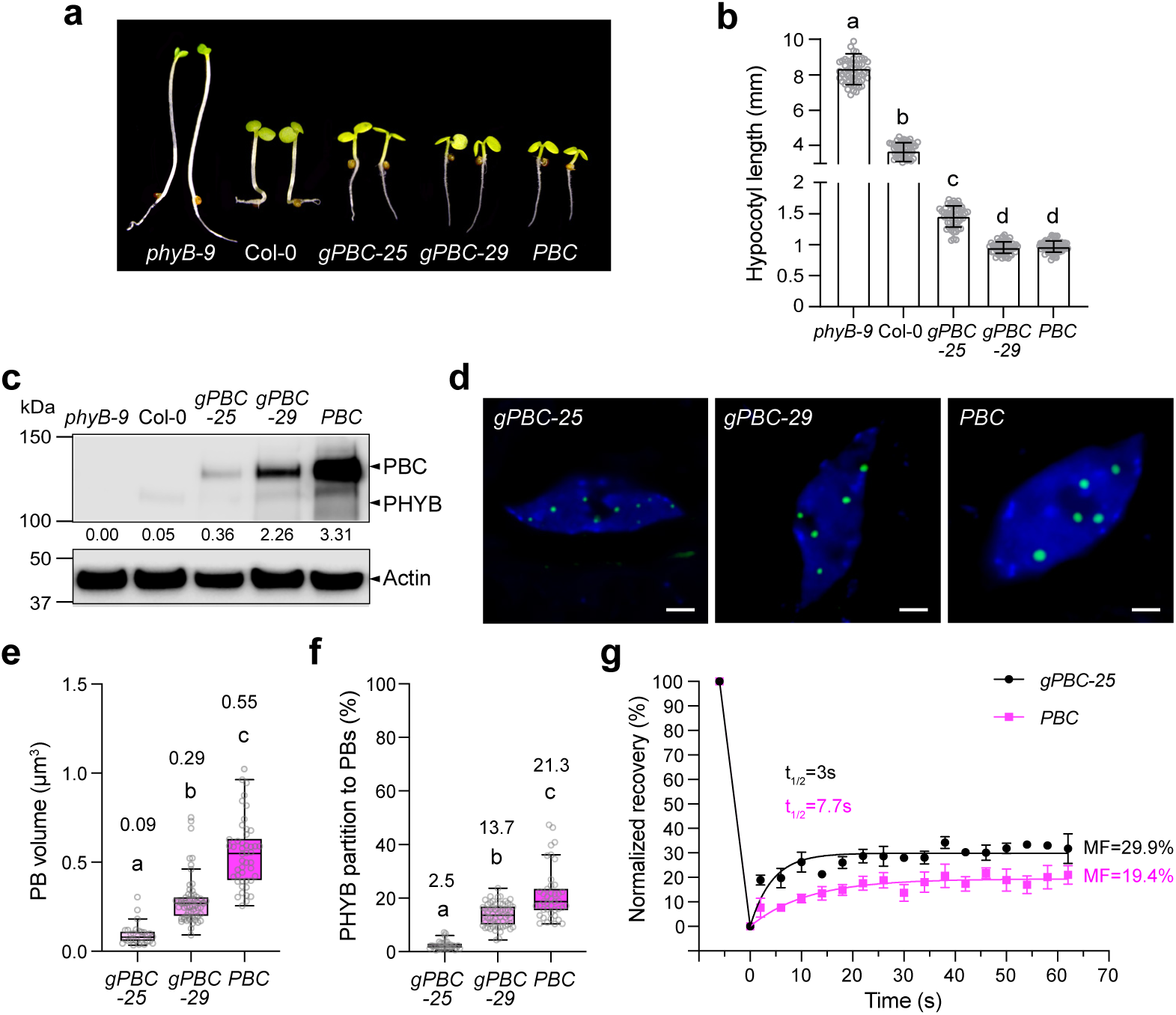
Increasing PHYB abundance alters PB size and dynamics. **a** Images of 4-d-old *phyB-9,* Col-0, *gPBC-25, gPBC-29* and *PBC* seedlings grown under 10 μmol m^−2^ s^−1^ R light at 21°C. **b** Hypocotyl length measurements of the seedlings shown in **a**. Error bars represent the s.d. (n > 50 seedlings); the centers of the error bars indicate the mean. Different letters denote statistically significant differences in hypocotyl length (ANOVA, Tukey’s HSD, *p*L ≤L 0.05). **c** Immunoblots showing the PHYB levels in the seedlings described in **a**. Actin was used as a loading control. The relative endogenous PHYB and PHYB-CFP levels normalized to the corresponding levels of actin are shown. The immunoblot experiments were independently repeated three times with similar results. **d** Confocal images showing representative PHYB-CFP PB patterns in cotyledon pavement epidermal cells from 4-d-old *gPBC-25, gPBC-29* and *PBC* seedlings grown under 10 μmol m^−2^ s^−1^ R light at 21°C. Scale bars are equal to 2Lµm. **e** Quantification of the PB volume in cotyledon epidermal cells from *gPBC-25, gPBC-29* and *PBC* as shown in **d**. Different letters denote statistically significant differences in volume (ANOVA, Tukey’s HSD, *p*L ≤L 0.05). **f** Quantification of the percentage of PHYB partitioned to PBs based on the total PHYB signal within the nuclei of *gPBC-25*, *gPBC-29* and *PBC* as shown in **d**. Box and whisker plots showing the percent of PHYB partitioned to PBs per nucleus. Different letters denote statistically significant differences in percentage (ANOVA, Tukey’s HSD, *p*L ≤L 0.05). For **e** and **f**, the numbers indicate the mean values. In the box and whisker plots, the boxes represent the 25th to 75th percentiles, and the bars are equal to the mean. **g** Results of FRAP experiments showing normalized fluorescence recovery plotted over time for PHYB-CFP in PBs of pavement cell nuclei from 4-d-old *gPBC-25* (black) and *PBC* (magenta) seedlings grown under 50 μmol m^−2^ s^−1^ R light. Values represent the mean, and error bars represent the s.e. of the mean of three biological replicates. The solid lines represent an exponential fit of the data. MF: mobile fraction; t_1/2_ represents the half-time of fluorescence recovery. The source data underlying the hypocotyl measurements in **b**, the immunoblots in **c**, and the PB characterization in **e**-**g** are provided in the Source Data file.

To test whether increasing the PHYB level enhances the size of PBs, we measured the volumes of PHYB-CFP PBs in *gPBC-25*, *gPBC-29*, and *PBC*. Indeed, the PB size increased with increases in the PHYB-CFP level (Fig. 1d,e). *PBC* had the largest PBs, which were on average more than 5-fold larger than those in *gPBC-25*, and the average PB volume of *gPBC-29* was three times larger than that of *gPBC-25* (Fig. 1e). The fraction of PHYB-CFP localized to PBs also increased with the PHYB-CFP level (Fig. 1f). These results support the predictions of the PHYB LLPS model that increasing PHYB abundance enlarges PBs, although these data did not assess whether the concentration of PHYB-CFP remained the same in the PBs and the surrounding nucleoplasm. We used fluorescence recovery after photobleaching (FRAP) to assess the exchange of PHYB-CFP molecules between PBs and the surrounding nucleoplasm. To our surprise, the PBs from *gPBC-25* and *PBC* exhibited significant differences in the dynamics of PHYB-CFP. The fluorescence recovery in *gPBC-25* was only 29.9%, suggesting that the majority of PHYB-CFP molecules were not mobile (Fig. 1g). The percentage of fluorescence recovery in *PBC* was further reduced to 19.9%, indicating that increasing PHYB-CFP abundance in *PBC* decreased the mobile fraction of PHYB-CFP in PBs, likely due to a transition to a gel-like state (Fig. 1g). Supporting this idea, PHYB-CFP in the PBs from *PBC* showed a decrease in the fluorescence recovery kinetics compared with *gPBC-25*, indicating a reduction in the diffusion rate of *PHYB-CFP* in *PBC* (Fig. 1g). Together, these results demonstrate that increasing PHYB abundance enhances PB size by recruiting a larger fraction of PHYB to the PB compartment and also induces a transition of PBs to a gel-like state that could retain PHYB for a longer time in PBs. Thus, this panel of lines with various PB sizes and PHYB-CFP dynamics provides a new opportunity to interrogate the function of PBs in PIF degradation.

### PIF5 is a short-lived protein localized in PBs

To investigate the function of PBs in PIF regulation, we turned to PIF5 as a model because despite PHYB-mediated PIF5 degradation^44,45,47^, PIF5 can accumulate in the light^36,45,47,54,55^, which could potentially allow us to monitor its subnuclear localization in the light. To that end, we generated transgenic lines expressing PIF5 fused with Myc and mCherry to the N-terminus (mCherry-PIF5) under the native *PIF5* promoter in the null *pif5-3* mutant background. Two independent transgenic lines with a single insertion of the transgene, named *mCherry-PIF5/pif5-3 #8* and *#9*, were selected for further analysis. The *pif5-3* mutant exhibited a short hypocotyl phenotype at 16°C. Because PIF5 is required for the warm temperature-induced hypocotyl elongation, the defect of *pif5-3* in hypocotyl growth became more pronounced at 27°C (Fig. 2a,b)^54^. The *mCherry-PIF5/pif5-3* lines rescued the hypocotyl-growth defects of *pif5-3* at both temperatures (Fig. 2a,b), indicating that mCherry-PIF5 was functional. Similar to endogenous PIF5, mCherry-PIF5 accumulated in the light and could be detected by immunoblotting using anti-PIF5 antibodies (Fig. 2c). Despite being controlled by the native *PIF5* promoter, the steady-state levels of mCherry-PIF5 in the transgenic lines were higher than that of endogenous PIF5 in Col-0. The mCherry-PIF5 level in line *#8* was more than 5-fold that of line #9 and more than 10-fold that of endogenous PIF5 in Col-0 (Fig. 2c). However, to our surprise, no mCherry signal could be detected by confocal microscopy in either of the *mCherry-PIF5/pif5-3* transgenic lines. We reasoned that this discrepancy might be attributable to a faster turnover rate of mCherry-PIF5 compared with the maturation time required for newly synthesized mCherry to become fluorescent. The reported maturation time of mCherry is more than 60 min^66^. If the half-life of mCherry-PIF5 is significantly shorter than 60 min, the newly synthesized mCherry-PIF5 would have been degraded before becoming fluorescent. If this were the case, the non-fluorescent mCherry-PIF5 protein should be detectable by immunolocalization. To test the hypothesis, we performed immunofluorescence staining using anti-Myc antibodies. Indeed, Myc-tagged mCherry-PIF5 could be detected by immunostaining. More interestingly, mCherry-PIF5 was localized to discrete punctate structures at both 16°C and 27°C (Fig. 2d). Simultaneously labeling endogenous PHYB using anti-PHYB antibodies demonstrated that mCherry-PIF5 colocalized with PHYB in PBs (Fig. 2d,e). Thus, these results indicate that mCherry-PIF5 is a short-lived protein that colocalizes with PHYB in PBs under the physiological PHYB concentration.

**Fig. 2.**
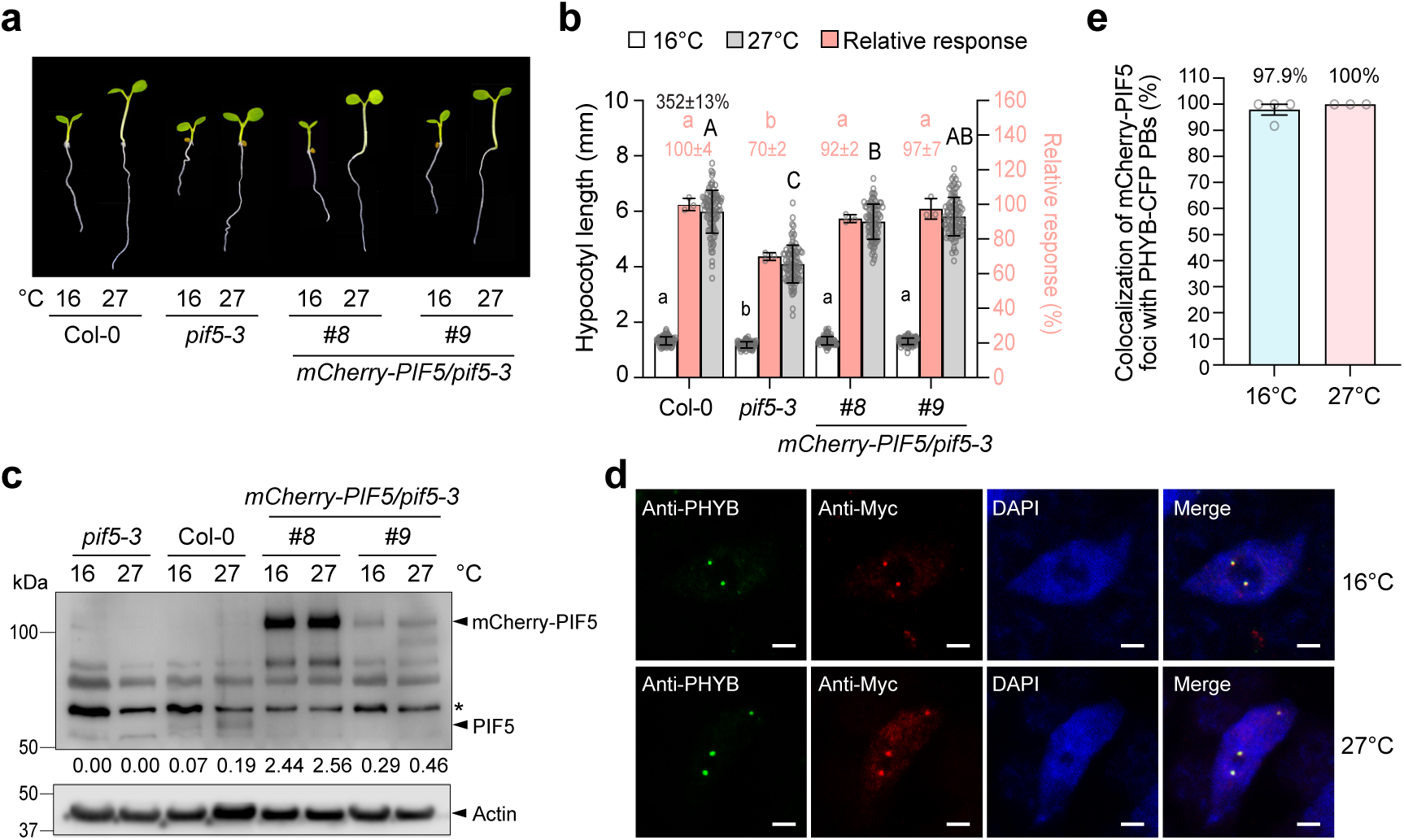
mCherry-PIF5 is a short-lived protein localized in PBs. **a** Images of 4-d-old seedlings of Col-0, *pif5-3* and two *mCherry-PIF5/pif5-3* (#*8* and #*9*) lines grown under 50 μmol m^−2^ s^−1^ R light at 16°C or 27°C. **b** Hypocotyl length measurements of the seedlings described in **a**. The open and gray bars represent hypocotyl length measurements at 16°C and 27°C, respectively. Error bars for the hypocotyl measurements represent the s.d. (n = 90 seedlings). Different lowercase and uppercase letters denote statistically significant differences in hypocotyl lengths at 16°C or 27°C, respectively (ANOVA, Tukey’s HSD, *p*L<L0.05, nL=L90 seedlings). The black number above the Col-0 data represents the percent increase in hypocotyl length at 27°C compared to 16°C (mean ± s.d., n = 90 seedlings). The pink bars show the relative response, which is defined as the hypocotyl response to 27°C of a mutant or transgenic line relative to that of Col-0 (set at 100%). Error bars for the relative responses represent the s.e. of three biological replicates. The centers of all error bars indicate the mean. Pink numbers show the mean ± s.e. of the relative responses. Different pink letters denote statistically significant differences in the relative responses (ANOVA, Tukey’s HSD, *p*L<L0.05, nL=L3 biological replicates). **c** Immunoblots showing the steady-state levels of PIF5 and mCherry-PIF5 in 4-d-old Col-0, *pif5-3*, and the *mCherry-PIF5* lines (*#8*, *#9*) grown under 50 μmol m^−2^ s^−1^ R light at 16°C or 27°C. PIF5 and mCherry-PIF5 were detected using anti-PIF5 antibodies. Actin was used as a loading control. The relative levels of PIF5 or mCherry-PIF5 normalized to actin are shown. The asterisk indicates a nonspecific band. **d** Confocal images showing the colocalization of mCherry-PIF5 and PHYB in PBs in cotyledon pavement epidermal nuclei from 4-d-old *mCherry-PIF5* line *#8* seedlings grown under 50 μmol m^−2^ s^−1^ R light at 16°C or 27°C. PHYB (green) and Myc-tagged *mCherry-PIF5* (red) were labeled via immunofluorescence staining using anti-PHYB and anti-Myc antibodies, respectively. Nuclei were stained with DAPI (blue). Scale bars are equal to 2Lµm. **e** Quantification of the colocalization of mCherry-PIF5 foci with PHYB-CFP PBs in the experiments described in **d**. Error bars represent the s.e. of at least three biological replicates. The source data underlying the hypocotyl measurements in **b**, the immunoblots in **c**, and the PB characterization in **e** are provided in the Source Data file.

### PHYB signaling both degrades and stabilizes PIF5

The fact that mCherry-PIF5 was short-lived and localized in PBs might suggest that mCherry-PIF5 degradation occurs in PBs. However, it was equally possible that mCherry-PIF5 was degraded in the nucleoplasm and that because mCherry-PIF5 was exchanged between the nucleoplasm and PBs, both the nucleoplasmic and PB pools of mCherry-PIF5 were short-lived. One way to distinguish between these two possibilities would be to test whether and how perturbing PBs alters PIF5 degradation. If PIF5 were degraded in PBs, increasing the size of PBs should recruit more PIF5 to PBs for degradation and therefore would be expected to accelerate PIF5 degradation and reduce the steady-state level of PIF5. To test the hypothesis, we examined the endogenous levels of PIF5 in the panel of lines expressing various amounts of PHYB (Fig. 1a-c). To our surprise, PIF5 showed a biphasic response to increases in PHYB abundance. In the range of zero to moderate levels of PHYB in *phyB-9*, Col-0, and *gPBC-25*, the steady-state level of PIF5 decreased with the increases in PHYB (Fig. 3a-c), supporting the current model that PHYB promotes PIF5 degradation in the light. However, unexpectedly, further increasing PHYB abundance in *gPBC-29* and *PBC* enhanced PIF5 accumulation (Fig. 3a,b). Because despite the changes in the PHYB level, the transcript levels of *PIF5* remained the same in *phyB-9*, Col-0, *gPBC-25*, *gPBC-29*, and *PBC* (Fig. 3c), the changes in the protein level of PIF5 must be due to either PIF5 translation or degradation. We next examined the degradation kinetics of PIF5 in Col-0, *gPBC-25*, and *PBC* by treating the cell lysates with the translation inhibitor cycloheximide and then monitoring the disappearance of PIF5 over time. PIF5 was degraded at a much faster rate in *gPBC-25* than in Col-0 (Fig. 3d,e). However, surprisingly, PIF5 degradation was attenuated dramatically in *PBC*. Thus, the accumulation of PIF5 in *PBC* and *gPBC-29* was most likely due to the PHYB-mediated stabilization of PIF5. These results suggest that PHYB signaling exerts opposing roles in both degrading and stabilizing PIF5, the balance of which can be adjusted by altering the PHYB level. Because a major change from *gPBC-25* to *PBC* was an increase in the size of PBs (Fig. 1d,e), these results raised a new hypothesis that while PHYB promotes PIF5 degradation in the nucleoplasm, it stabilizes PIF5 in PBs. As such, increasing the PB size in *PBC* could switch the balance towards PIF5 stabilization by recruiting more PIF5 to PBs.

**Fig. 3.**
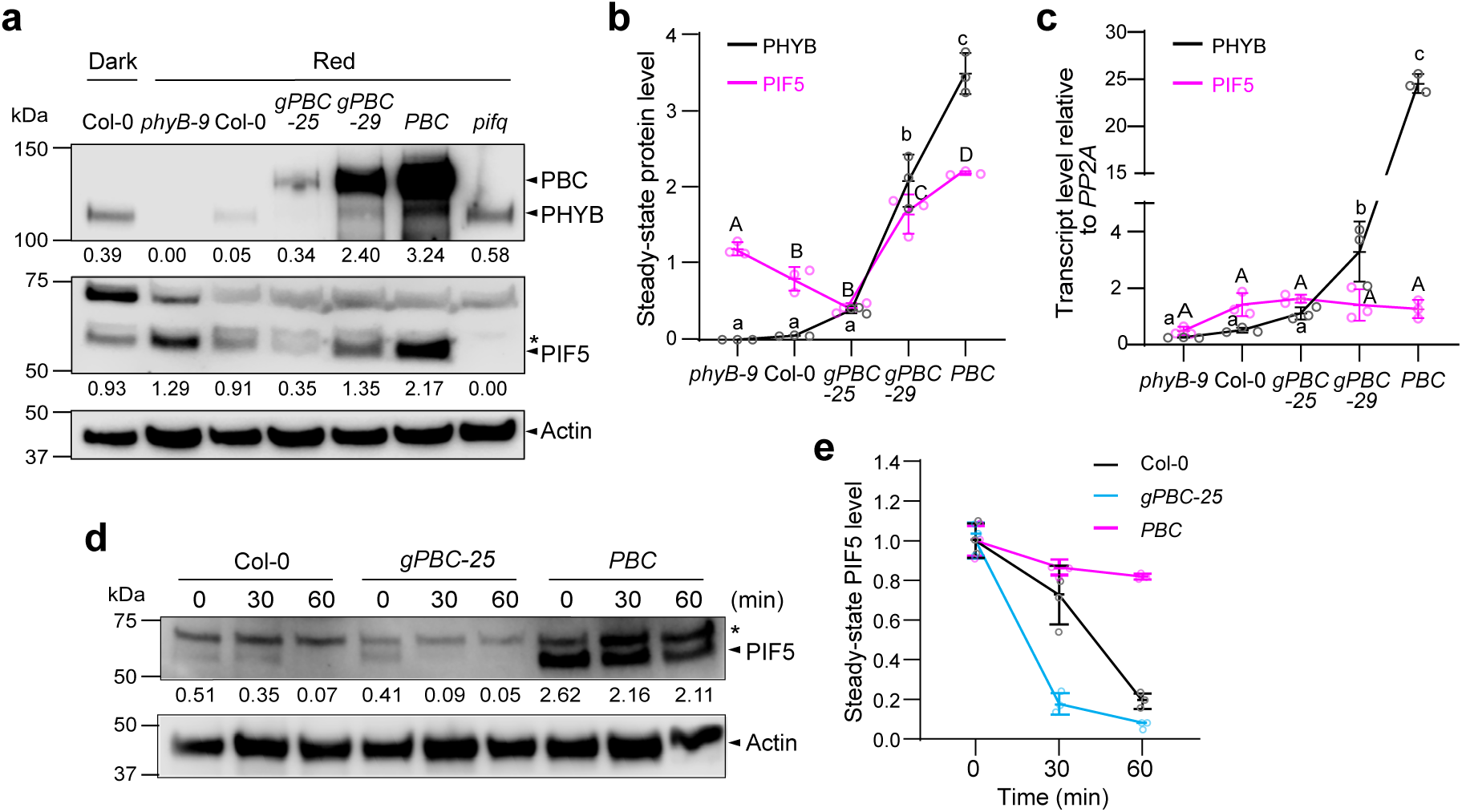
Increasing PHYB abundance provokes a biphasic PIF5 response. **a** Immunoblots showing PHYB and PIF5 levels in 4-d-old *phyB-9,* Col-0, *gPBC-25, gPBC-29* and *PBC* seedlings grown under 10 μmol m^−2^ s^−1^ R light at 21°C. 4-d-old dark-grown Col-0 and R-light-grown *pifq* were used as positive and negative controls for PIF5, respectively. The immunoblot experiments were independently repeated three times with similar results. **b** Quantification of the PHYB and PIF5 levels shown in **a**. Error bars represent the s.d. of three independent replicates. The centers of the error bars indicate the mean values. Different lowercase and uppercase letters denote statistically significant differences in the abundance of PHYB and PIF5, respectively (ANOVA, Tukey’s HSD, *p* < 0.05, n = 3 biological replicates). **c** Quantitative real-time PCR (qRT-PCR) analysis of the transcript levels of *PHYB* and *PIF5* in seedlings shown in **a**. Different lowercase and uppercase letters denote statistically significant differences in *PHYB* and *PIF5* transcripts, respectively (ANOVA, Tukey’s HSD, *p* < 0.05, n = 3 biological replicates). **d** Immunoblots showing the degradation kinetics of PIF5 in Col-0, *gPBC-25,* and *PBC*. 4-d-old Col-0, *gPBC-25* and *PBC* seedlings were incubated with cycloheximide and collected at the indicated time points. Total proteins were extracted and subjected to immunoblotting analysis using anti-PIF5 antibodies. For **a** and **d**, actin was used as a loading control. The asterisk indicates a nonspecific band. The relative PHYB or PIF5 levels normalized to the corresponding levels of actin are shown. **e** Quantification of the relative PIF5 protein levels shown in **d**. Error bars represent the s.d. of three independent replicates. The source data underlying the immunoblots in **a**, **b**, **d**, and **e**, and the qRT-PCR analysis in **c** are provided in the Source Data file.

### PHYB recruits and stabilizes PIF5 in PBs

To further examine the role of PBs in stabilizing PIF5, we reasoned that if PIF5 were stabilized due to its enhanced recruitment to the enlarged PBs in *PBC*, we might be able to observe the accumulation of stabilized or longer-lived mCherry-PIF5 proteins in PBs in *PBC* via confocal microscopy. To that end, we generated transgenic lines expressing mCherry-PIF5 tagged with human influenza hemagglutinin (HA) (HA-mCherry-PIF5) under the constitutive *UBIQUITIN 10* promoter in *PBC* (*mCherry-PIF5/PBC*). We selected two *mCherry-PIF5/PBC* lines (*#1* and *#9*) for further analysis. The *mCherry-PIF5/PBC* lines were taller than *PBC* at both 16°C and 27°C, suggesting that the HA-tagged mCherry-PIF5 was functional (Fig. 4a,b). As endogenous PIF5, mCherry-PIF5 could accumulate in both *mCherry-PIF5/PBC* lines in the light at 16°C and 27°C (Fig. 4c). Similar to the *mCherry-PIF5/pif5-3* lines (Fig. 3c), the mCherry-PIF5 levels in the *mCherry-PIF5/PBC* lines were more than 10-fold greater than that of the endogenous PIF5 in Col-0 (Fig. 4c). However, unlike the *mCherry-PIF5/pif5-3* lines, we could observe the fluorescent signal of mCherry-PIF5 in the *mCherry-PIF5/PBC* lines using confocal microscopy (Fig. 4d). mCherry-PIF5 was also colocalized with PHYB-CFP in PBs in the *mCherry-PIF5/PBC* lines at both 16°C and 27°C in the light (Fig. 4d,e). In dark-grown *mCherry-PIF5/PBC* seedlings where there were no PHYB-CFP PBs, mCherry-PIF5 was evenly dispersed in the nucleoplasm (Fig. 4f), implying that mCherry-PIF5 was recruited to PBs in the light by PHYB. The fact that mCherry-PIF5 became detectable by confocal microscopy suggested that the half-life of the mCherry-PIF5 in *mCherry-PIF5/PBC* was significantly longer than that in *mCherry-PIF5/pif5-3*. Supporting this conclusion, like the enhanced accumulation of endogenous PIF5 in *PBC* compared with Col-0, mCherry-PIF5 accumulated to higher levels in *mCherry-PIF5/PBC* compared with *mCherry-PIF5/pif5-3* (Figs 3a,b and 4c). Moreover, the degradation kinetics of mCherry-PIF5 became slower compared with that in *mCherry-PIF5/pif5-3* (Fig. 4g,h). It is important to note that the degradation kinetics of mCherry-PIF5 (Fig. 4g,h) was considerably faster than that of endogenous PIF5 (Fig. 3d,e)^47^, this could be attributed to either the mCherry tag or the overexpression of mCherry-PIF5 and the resulted change in their PB/nucleoplasm partitioning. Together, these results support the model that PHYB recruits and stabilizes PIF5 in PBs and that the action of PHYB in PIF5 stabilization is enhanced in *PBC* by retaining more PIF5 in the enlarged PBs.

**Fig. 4.**
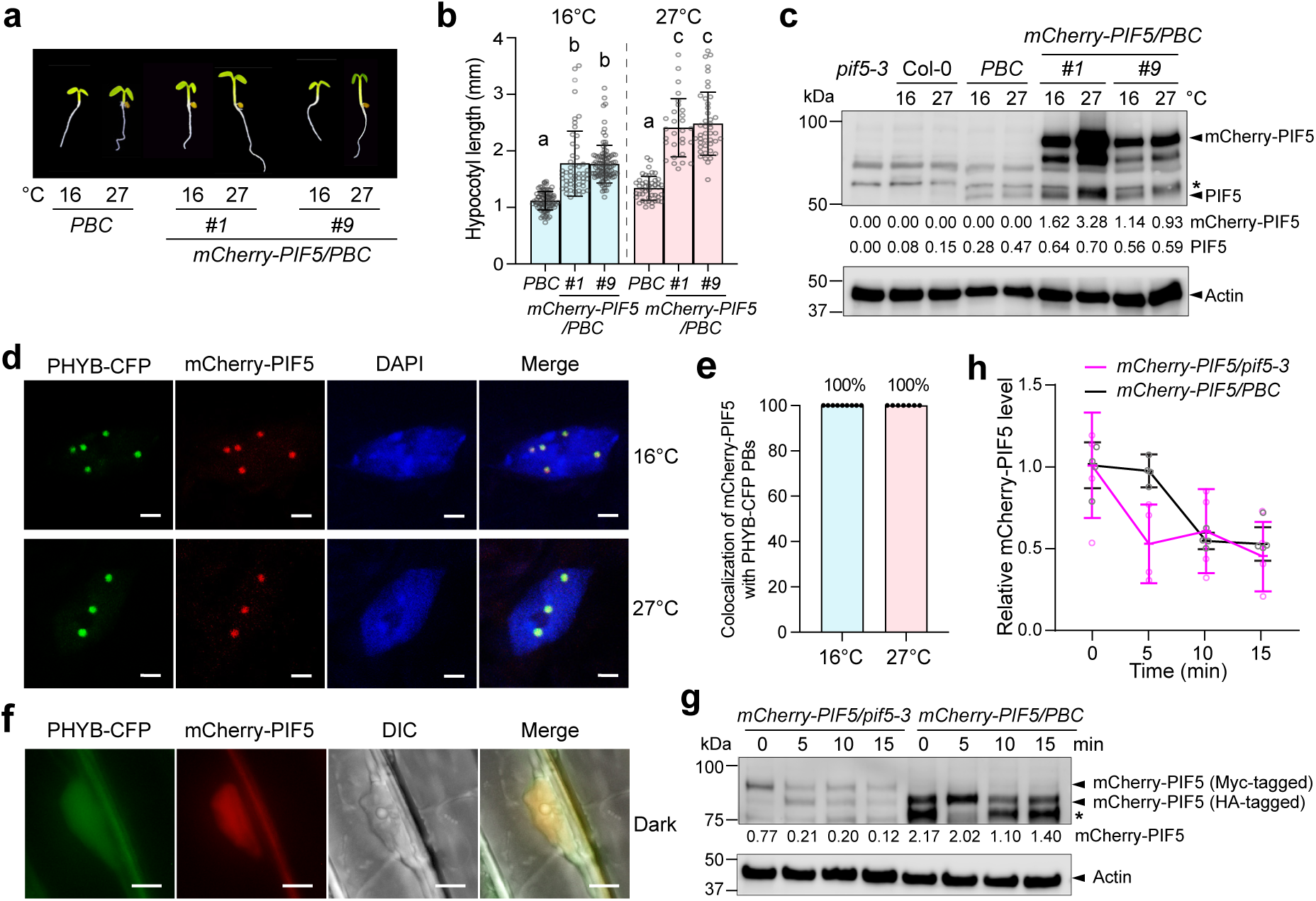
PHYB recruits and stabilizes PIF5 in PBs. **a** Images of 4-d-old *PBC* and *mCherry-PIF5/PBC* (#*1* and #*9*) seedlings grown under 10 μmol m^−2^ s^−1^ R light at either 16°C or 27°C. **b** Hypocotyl length measurements of the seedings described in **a**. Error bars represent the s.d., and the centers represent the mean (n ≥ 32 seedlings). Samples labeled with different letters exhibited statistically significant differences in hypocotyl length (ANOVA, Tukey’s HSD, *p* ≤L0.05). **c** Immunoblots showing the steady-state levels of PIF5 and mCherry-PIF5 in 4-d-old *pif5-3*, Col-0, *PBC*, and *mCherry-PIF5/PBC* (*#1*, *#9*) seedlings grown under 10 μmol m^−2^ s^−1^ R light at either 16°C or 27°C. **d** Confocal images showing the colocalization of mCherry-PIF5 with PHYB-CFP in PBs in cotyledon pavement cells from 4-d-old *mCherry-PIF5/PBC #9* seedlings grown under 10 μmol m^−2^ s^−1^ R light at either 16°C or 27°C. Scale bars are equal to 2Lµm. **e** Quantification of the colocalization of mCherry-PIF5 foci with PHYB-CFP PBs in the experiments described in **d**. Error bars represent the s.e. (n = 9 biological replicates for the 16°C samples, n = 7 biological replicates for the 27°C samples), and the centers represent the mean. **f** Fluorescence microscope images showing the localization of PHYB-CFP and mCherry-PIF5 in hypocotyl epidermal cells from 4-d-old dark-grown *mCherry-PIF5/PBC #9* seedings. The DIC image shows the nucleus. Scale bars are equal to 5 µm. **g** Immunoblots showing the degradation kinetics of Myc-tagged mCherry-PIF5 in *mCherry-PIF5/pif5-3* and HA-tagged mCherry-PIF5 in *mCherry-PIF5/PBC*. 4-d-old seedlings of *mCherry-PIF5/pif5-3 #8* and *mCherry-PIF5/PBC #9* were incubated with cycloheximide and collected at the indicated time points. For **c** and **g**, PIF5 and mCherry-PIF5 were detected using anti-PIF5 antibodies. Actin was used as a loading control. The asterisk indicates a nonspecific band. The relative levels of PIF5 and mCherry-PIF5, normalized to the corresponding levels of actin, are shown below each lane. **h** Quantification of the relative mCherry-PIF5 levels shown in **g**. Error bars represent the s.d. of four independent replicates. The source data underlying the hypocotyl measurements in **b**, the immunoblots in **c, g**, and **h**, and the quantification of the colocalization of mCherry-PIF5 and PHYB-CFP in **e** are provided in the Source Data file.

### Reducing PB size accelerates PIF5 degradation

If PB formation phase-separates the opposing PHYB signaling actions of PIF5 degradation and stabilization, altering PB size and therefore the PB/nucleoplasm partitioning of PHYB and PIF5 should shift the balance of the two antagonistic PHYB actions and result in a change in the level of PIF5 and its signaling output. For example, if the above model is correct, because recruiting PIF5 to PBs prevents PIF5 degradation in the surrounding nucleoplasm, reducing PB size in *mCherry-PIF5/PBC* under dimmer light should increase the nucleoplasmic fractions of PHYB and PIF5 and therefore rebalance PHYB signaling from stabilizing PIF5 to degrading PIF5. To test this hypothesis, we compared the size and dynamics of PBs as well as PIF5 degradation in *mCherry-PIF5/PBC* #9 grown under 10 μmol m^−2^ s^−1^ or dim 0.5 μmol m^−2^ s^−1^ R light. As expected, the dimmer R light reduced PHYB activity and therefore promoted hypocotyl elongation in Col-0 (Fig. 5a,b). The *mCherry-PIF5/PBC* seedlings grown under 0.5 μmol m^−2^ s^−1^ R light were also significantly taller than the *mCherry-PIF5/PBC* seedlings grown under 10 μmol m^−2^ s^−1^ and *PBC* seedlings grown 0.5 μmol m^−2^ s^−1^ R light (Fig. 5a,b), which suggests that mCherry-PIF5 in *mCherry-PIF5/PBC* was more abundant and/or more active under the dim light. As expected, PHYB-CFP PBs became significantly smaller under the dim light (Fig. 5c,d)^28^. Interestingly, mCherry-PIF5 was detectable using confocal microscopy in only 75.3% of PBs in the dim light (Fig. 5e) as opposed to in 100% PBs under strong R light (Fig. 4e), suggesting counterintuitively that mCherry-PIF5 might be less stable in dim light. Supporting this conclusion, the steady-state levels of both mCherry-PIF5 and endogenous PIF5 decreased dramatically in *mCherry-PIF5/PBC* under 0.5 μmol m^−2^ s^−1^ R light compared to 10 μmol m^−2^ s^−1^ R light (Fig. 5f). The dimmer light treatment also reduced the PIF5 level in *PBC*, indicating that the low-light effect in *mCherry-PIF5/PBC* was not caused by the overexpression of mCherry-PIF5. Consistently, the degradation kinetics of mCherry-PIF5 were dramatically increased in *mCherry-PIF5/PBC* under dim R light compared to that under strong R light (Fig. 5g). PHYB-CFP became more dynamic in the small PBs under 0.5 μmol m^−2^ s^−1^ R light, suggesting that dimmer light promoted a gel-to-liquid transition (Fig. 5h). Intriguingly, the dynamics of mCherry-PIF5 stayed the same between high and low light conditions (Fig. 5i). These results suggest that although the large and small PBs might have different PIF5-binding capacities, their PIF5 binding affinities were similar. Therefore, reducing the PB size promotes PIF5 degradation likely by enhancing the partitioning of PHYB and PIF5 to the nucleoplasm compartment. Together, these results provide further evidence supporting our conclusion that recruiting PIF5 to PBs stabilizes PIF5 by preventing its degradation in the surrounding nucleoplasm.

**Fig. 5.**
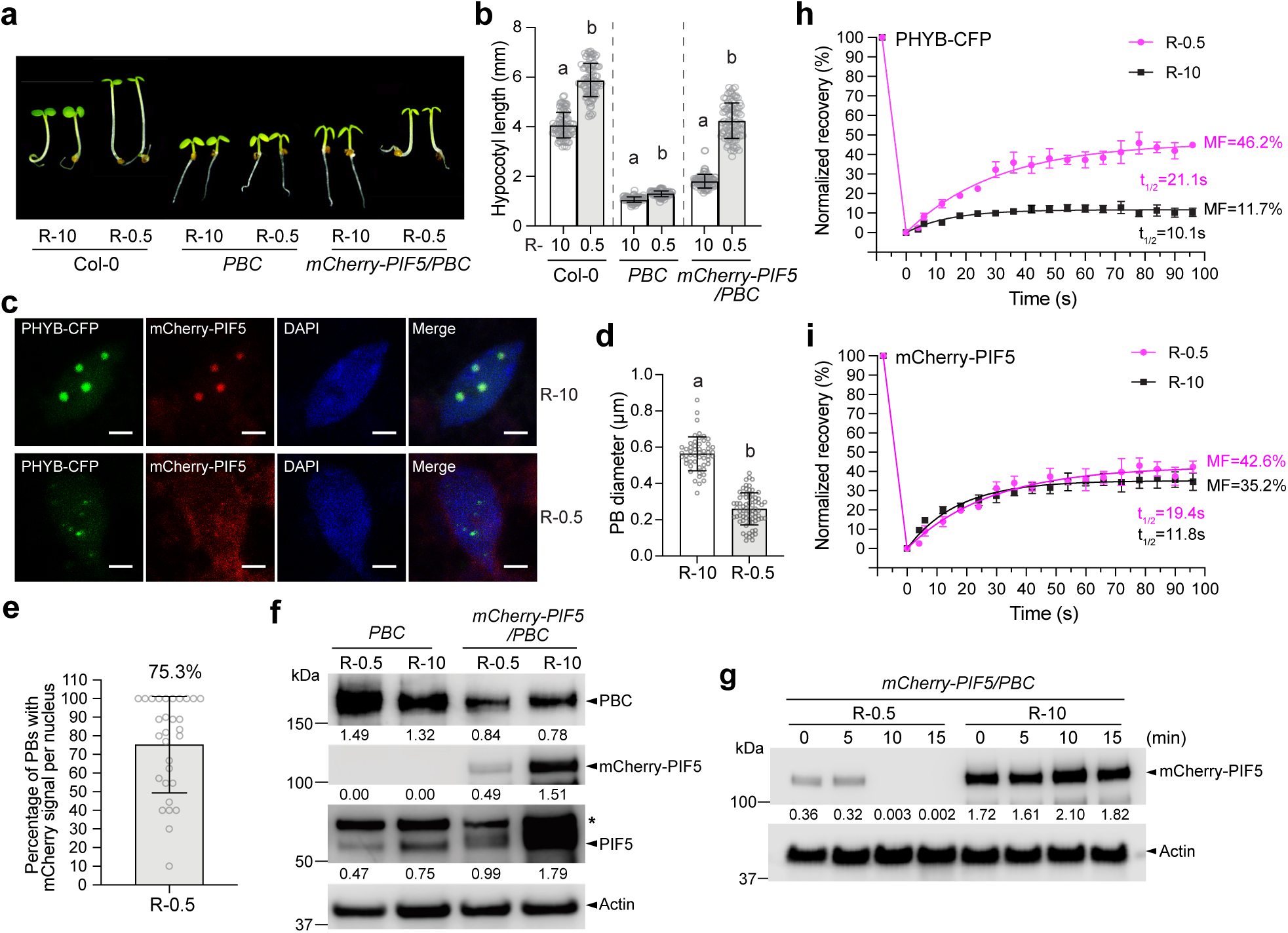
Reducing PB size accelerates PIF5 degradation. **a** Images of 4-d-old Col-0, *PBC* and *mCherry-PIF5/PBC #9* seedlings grown under either 10 μmol m^−2^ s^−1^ or 0.5 μmol m^−2^ s^−1^ R light (R-10 and R-0.5, respectively). **b** Hypocotyl length measurement of the seedlings described in **a**. Error bars represent the s.d., and the centers represent the mean (n = 90 seedlings). Samples labeled with different letters exhibited statistically significant differences in hypocotyl length (Student’s t-test, *p* ≤L0.05). **c** Confocal images showing the PB patterns in cotyledon pavement epidermal cells from the *mCherry-PIF5/PBC* seedlings described in **a**. Scale bars are equal to 2Lµm. **d** Quantification of the sizes of the PBs described in **c**. Error bars represent the s.d., and the center represents the mean (n = 53 PBs for the R-10 samples, n = 72 for the R-0.5 samples). Different letters denote statistically significant differences in PB size (Student’s t-test, *p* ≤ 0.05). **e** Quantification of the percentage PHYB-CFP PBs with detectable mCherry-PIF5 fluorescence in *mCherry-PIF5/PBC* seedlings grown under 0.5 μmol m^−2^ s^−1^ R light. The error bar represents the s.d. and the center represents the mean (n = 30). **f** Immunoblots of PHYB-CFP, endogenous PIF5 and mCherry-PIF5 in 4-d-old *PBC* and *mCherry-PIF5/PBC #9* seedlings grown under either 10 μmol m^−2^ s^−1^ or 0.5 μmol m^−2^ s^−1^ R light. PHYB-CFP was detected using anti-PHYB antibodies, and mCherry-PIF5 and PIF5 were detected using anti-PIF5 antibodies. Actin was used as a control. The relative levels of PHYB-CFP, endogenous PIF5 and mCherry-PIF5, normalized to the corresponding actin levels, are shown under each lane. The asterisk indicates a nonspecific band. **g** Immunoblots showing the degradation kinetics of mCherry-PIF5 in 4-d-old *mCherry-PIF5/PBC #9* seedlings grown under either 10 μmol m^−2^ s^−1^ or 0.5 μmol m^−2^ s^−1^ R light. Seedlings were incubated with cycloheximide and collected at the indicated time points. Total proteins were extracted and subjected to immunoblotting analysis using anti-PIF5 antibodies. Actin was used as a loading control. The relative mCherry-PIF5 levels normalized to the corresponding levels of actin are shown. **h** Results of FRAP experiments showing normalized fluorescence recovery plotted over time for PHYB-CFP in the *mCherry-PIF5/PBC #9* seedlings described in **a**. **i** Results of FRAP experiments showing normalized fluorescence recovery plotted over time for mCherry-PIF5 in the *mCherry-PIF5/PBC #9* seedlings described in **a**. For **h** and **i**, values represent the mean, and error bars represent the s.e. of at least three biological replicates. Solid lines represent the exponential fit of the data. MF: mobile fraction. t_1/2_ represents the half-time of fluorescence recovery. The source data underlying the hypocotyl measurements in **b**, the PB characterization in **d** and **e**, the immunoblots in **f** and **g**, and the FRAP results in **h** and **i** are provided in the Source Data file.

## DISCUSSION

Reports published more than 20 years ago described how light triggers the localization of photoactivated PHYB to discrete PBs, creating two spatially separated pools of PHYB—a concentrated pool in PBs and a dilute pool in the surrounding nucleoplasm—and that PB size depends on the amount of active PHYB and therefore could be directly regulated by light intensity and quality (i.e., the R/FR ratio)^22,23,28^. However, the functional significance of this spatial segregation of active PHYB remained frustratingly elusive. Here, using PIF5 as a model, we show that PHYB recruits PIF5 as a client to PBs, and surprisingly, that PHYB exerts two opposing functions in both degrading and stabilizing PIF5. Different from previous studies on PBs, we perturbed PBs by increasing PHYB abundance. This approach allowed us to uncouple the function of PHYB in PBs from that in the surrounding nucleoplasm. Our results support the model that the condensation of PHYB phase-separates the opposing PHYB signaling action of PIF5 degradation and stabilization into two subnuclear compartments: a PIF5-stabilizing environment in PBs and a PIF5-degrading environment in the surrounding nucleoplasm (Fig. 6a). As such, environmentally controlled PB dynamics can regulate the PB/nucleoplasmic partitioning of PHYB, PIF5, and other signaling components, thereby enabling an environmentally sensitive counterbalancing mechanism for titrating the nucleoplasmic concentration of PIF5 and the signaling output of PHYB (Fig. 6b). Because control of the stability of PHYB-associated signaling components is a major mechanism of light signaling^18,19,41^, the novel PB-dependent counterbalancing mechanism provides the framework for regulating a plethora of PHYB-interacting signaling molecules in diverse plant environmental responses. We propose that this PB function represents a general function of biomolecular condensates that allows distinct variations of a cellular process or signaling pathway to coexist and interact in order to generate dynamically adjustable integrated outputs within a single subcellular space.

**Fig. 6.**
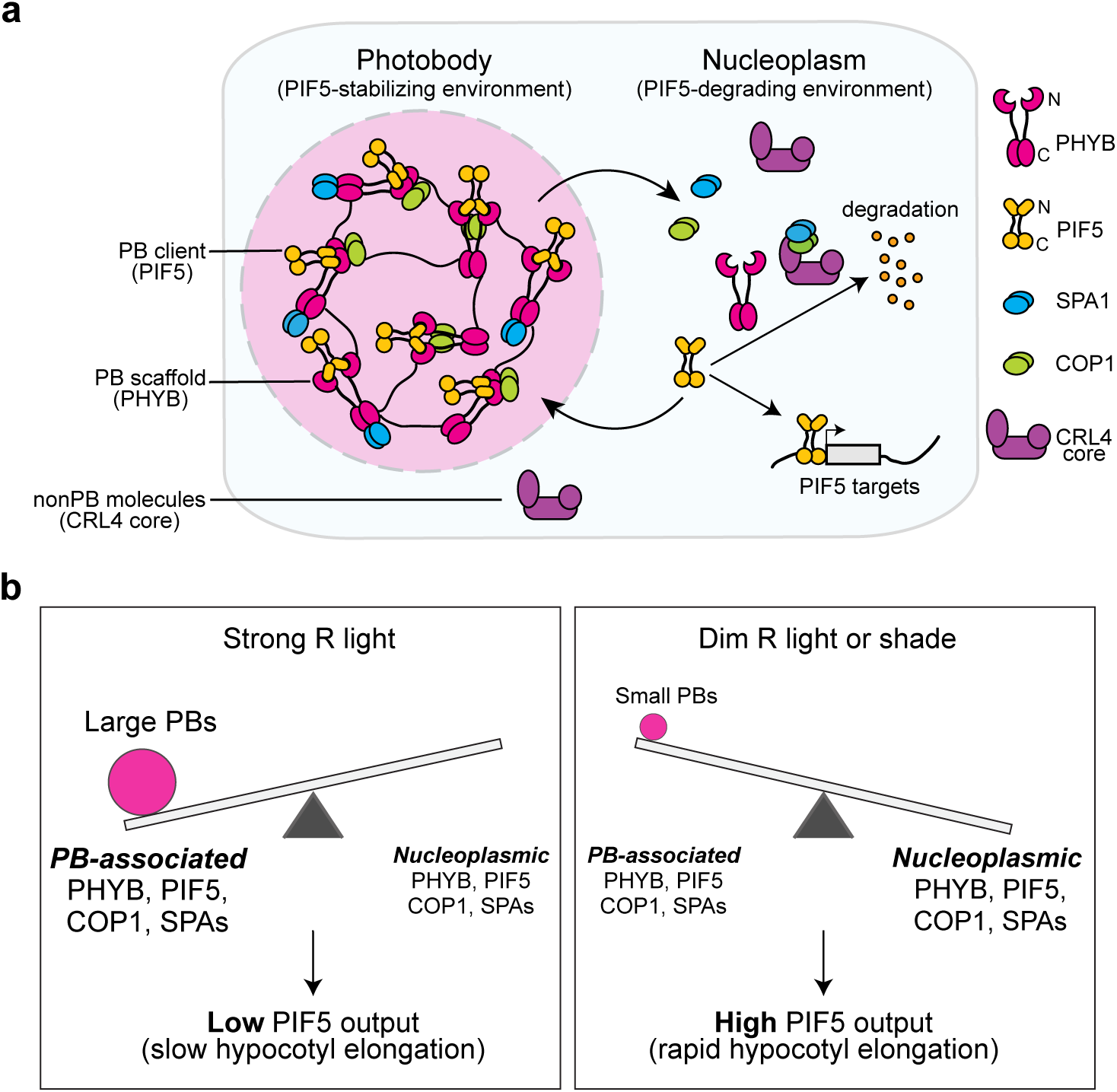
A PB-enabled counterbalancing model for titrating PHYB-mediated environmental responses. **a** Schematic illustration of the function of PBs in phase-separating the opposing PHYB signaling actions of PIF5 degradation and stabilization. The condensation of PHYB creates a PIF5-stabilizing environment in PBs to counteract PIF5 degradation in the surrounding nucleoplasm. In the nucleoplasm, PHYB promotes PIF5 degradation by triggering its phosphorylation and subsequent ubiquitylation by the CRL4^COP1/SPA^ E3 ubiquitin ligases^44,45,47^. PHYB serves as a scaffold component to recruit PIF5 as a client to PBs via direct interaction. PHYB stabilizes PIF5 by selectively recruiting COP1 and SPAs, but not the CRL4 core, to PBs^26,58,72^. The high concentration of PHYB in PBs disrupts the COP1-SPA complex via distinct interactions of COP1 with PHYB’s N-terminal photosensory module and SPAs with PHYB’s C-terminal output module^51–53,69,71^. **b** Model for an environmentally-sensitive PB-enabled counterbalancing mechanism to titrate nucleoplasmic PIF5 and environmental responses. Changes in the intensity and composition of light (and also in temperature) directly control the amount of active PHYB and thus the size of PBs to regulate the PB-to-nucleoplasm partitioning of PHYB, PIF5 and other PB constituents, thereby titrating the nucleoplasmic PIF5 and its signaling output. Strong R light increases the amount of active PHYB and enlarges PBs, thereby sequestering a greater fraction of PHYB and PIF5 in PBs, which stabilizes PIF5 and simultaneously reduces the nucleoplasmic PIF5 and its signaling output. This mechanism allows hypocotyl to grow slowly in the light. Dim R light or shade reduces the amount of active PHYB and PB size, thereby enhancing the nucleoplasmic fraction of PHYB and PIF5, which simultaneously promotes PIF5 degradation and its transcriptional output. The latter mechanism accelerates hypocotyl elongation.

Our results indicate that PHYB recruits PIF5 to PBs for PIF5 stabilization. We demonstrated for the first time that PIF5 localizes to PBs. This conclusion is supported by the colocalization of mCherry-PIF5 with endogenous PHYB in PBs in *mCherry-PIF5/pif5-3* and with PHYB-CFP PBs in *mCherry-PIF5/PBC* (Figs. 2d,e and 4d,e). We showed that the slow maturation time of FPs poses a major obstacle in observing FP-tagged PIFs in live cells. The short-lived mCherry-PIF5 in the *mCherry-PIF5/pif5-3* lines could only be detected using immunofluorescence staining, not via confocal microscopy (Fig. 2d). This technical hurdle might explain the surprisingly low number of detailed investigations into the subcellular localization of FP-tagged PIFs compared with the overwhelming number of reports on PIFs’ functions. Although the slow maturation of mCherry hindered our ability to observe mCherry-PIF5, we serendipitously found that the detectability of mCherry fluorescence could be used as an internal reporter to assess the stability or half-life of mCherry-PIF5 in live cells. When PIF5 degradation was attenuated in *mCherry-PIF5/PBC* (Fig. 4g), the fluorescence of mCherry-PIF5 became detectable using confocal microscopy (Fig. 4d). The fact that longer-lived mCherry-PIF5 in *mCherry-PIF5/PBC* was colocalized with PHYB-CFP in PBs provides direct evidence demonstrating that mCherry-PIF5 was retained and stabilized mCherry-PIF5 in PBs (Fig. 4d,e). Interestingly, mCherry-PIF5 in *mCherry-PIF5/PBC* localized to PBs only in the light but not in darkness (Fig. 4d,f), supporting the idea that PIF5 was recruited to PBs by PHYB, likely via direct interaction^44^. This conclusion corroborates the proteomics results that PBs comprise PHYB and its primary and secondary interacting signaling components^26,58^. PIF5 was not identified in the reported proteomics analysis of PB components, which may be due to the timing and conditions used in those studies as the expression of PIF5 was controlled by the circadian clock^67^. One concept of biomolecular condensates is the concept of scaffolds and clients^68^. Scaffold molecules are considered the drivers of phase separation, whereas molecules that are recruited into biomolecular condensates formed by scaffolds are called clients^68^. Applying this concept to PBs, the current data suggest that PHYB is a scaffold component, whereas PIF5 is a client component recruited to PBs, likely via direct interactions with PHYB and other PB components (Fig. 6a). Supporting this model, altering PB size by manipulating the concentration of active PHYB changed the dynamics of the scaffold PHYB but not that of the client (PIF5) (Fig. 5h,i), suggesting that changing PB size altered only PB’s binding capacity but not affinity to PIF5. Together, our results reveal that in addition to promoting PIF5 degradation, PHYB also stabilizes PIF5 by recruiting PIF5 to PBs, providing a mechanism that allows PIF5 to accumulate in the light.

PHYB-mediated degradation of PIFs has been invoked as the central mechanism of light signaling^18,41,42^. PIF5 is relatively stable in dark-grown seedlings and, upon exposure to light, PHYA and PHYB induce the rapid phosphorylation and degradation of PIF5^44,45,47^. However, the site of PIF5 degradation in the nucleus had not been previously studied. Our results support the model that PHYB condensation phase-separates the PIF5-stabilizing environment in PBs from the PIF5-degrading environment in the surrounding nucleoplasm (Fig. 6a). The mechanism that distinguishes the PIF5-stabilizing environment in PBs and the PIF5-degrading environment in the surrounding nucleoplasm requires further investigation. Previous studies have suggested that PIF5 degradation is mediated by the CRL4^COP1/SPA^ E3 ubiquitin ligase^47^. PHYB interacts directly with COP1 and SPAs via different domains: while the N-terminal photosensory module interacts with COP1^69^, the C-terminal output module confers a strong interaction with SPAs^51,52,70^. As such, PHYB inhibits the activity of CRL4^COP1/SPA^ by blocking the COP1-SPA interaction^51–53,69,71^. Both COP1 and SPAs have been identified as PB components^26,58^. It is likely that the high concentration of PHYB in PBs facilitates the dissociation of COP1-SPA complexes (Fig. 6a). In addition, cullin4 was not identified as a PB constituent^26,59^ and instead was shown to disperse evenly in the nucleoplasm^72^. Therefore, it is possible that the functional CRL4^COP1/SPA^ complex could be assembled only in the nucleoplasm, providing another mechanism for the distinct functions between PBs and the surrounding nucleoplasm (Fig. 6a). In this model, the dynamic changes in PBs would also regulate the amount of functional CRL4^COP1/SPA^ complex in the nucleoplasm.

Our results indicate that PHYB condensation enables an environmentally sensitive counterbalancing mechanism to titrate environmental responses (Fig 6b). Changes in light intensity and composition (and also temperature) directly control the amount of active PHYB and thus the size of PBs to regulate the PB-to-nucleoplasmic partitioning of PHYB, PIF5 and other PB constituents, thereby titrating the nucleoplasmic PIF5 and its signaling output (Fig. 6b). Supporting this model, PB size correlated positively with PIF5 stabilization and negatively with PIF5 degradation. A transition from small PBs in *mCherry-PIF5/pif5-3* to large PBs in *mCherry-PIF5/PBC* switched the balance of PHYB signaling from degrading to stabilizing PIF5 (Figs. 2-4). Conversely, transferring *mCherry-PIF5/PBC* seedlings to dim light, which induced the transition from large PBs to small PBs, accelerated PIF5 degradation (Fig. 5). The current data suggest that localization of PIF5 to PBs impedes its functions in promoting hypocotyl elongation. This model corroborates the proposed function of PBs for sequestering PIF7^73,74^. Enhancing the nucleoplasmic pool of mCherry-PIF5 in dim light promoted hypocotyl elongation (Fig. 5a,b). However, counterintuitively, the levels of both endogenous PIF5 and mCherry-PIF5 in *mCherry-PIF5/PBC* were reduced in the dim light compared to those in the strong light condition (Fig. 5f), indicating that the steady-state level of PIF5 does not correlate with its function in promoting hypocotyl elongation, likely due to enhanced degradation in the nucleoplasm. Another possibility is that PIF5 degradation is associated or coupled with its transcription activation activity. A similar mechanism was proposed for PIF3 as blocking PIF3 degradation in the *hmr* mutant attenuates the activation of its target genes^62^.

The function of PBs in stabilizing PIF5 may not be simply extrapolated to other PIFs as PIFs are degraded via distinct mechanisms. The degradation of PIF1 is mediated by CRL4^COP1/SPA^ and CRL1^CTG10^ with the F-box protein COLD TEMPERATURE GERMINATION 10 (CTG10) as the substrate recognition subunit^70,75^, whereas PIF3 is degraded by CRL3^LRB^ and CRL1^EBF^. PBs were proposed to be required for PIF1 and PIF3 degradation^7,24,29,60,61,63,64^. However, it remains unclear whether PIF3 is degraded in PBs. Although PIF4 can also accumulate in the light^36,45,54^, PIF4 is ubiquitylated by the cullin3-based E3 ligase CRL3^BOP1/2^ using BLADE ON PETIOLE (BOP1/2) as the substrate recognition components^76^. When coexpressed with PHYB, FP-tagged PIF4, as well as PIF3 and PIF6/PIL2, localized with PHYB in biomolecular condensates in mammalian cells or protoplasts in a PHYB-dependent manner, supporting the idea that PHYB may be able to recruit PIF4 to PBs^37–39^. However, the PB localization of PIF4 in *Arabidopsis* has not been carefully studied, and the potential role of PBs in stabilizing PIF4 in the light remains to be experimentally verified.

A major difference that distinguishes this study from the previous investigations about PBs is the approach used to perturb PBs. Because components can diffuse between membraneless organelles and the surrounding environment, a major challenge in studying the function of nuclear bodies has been the difficulty in dissecting the functional role of components in nuclear bodies vs the surrounding nucleoplasm. Conventional approaches using loss-of-function mutants of nuclear body components may yield a correlation between assembly and a functional output; however, such a correlation is usually not sufficient to discern the function of nuclear bodies from functions in the surrounding nucleoplasm. Here, we perturbed the PB size by increasing PHYB abundance. If PB formation is driven by the LLPS of PHYB, increasing the PHYB level is expected to only enlarge the PB size without changing the concentrations of PHYB in the nucleoplasm and therefore may specifically enhance the functional output within PBs^4,65^. Consistent with the hypothesis, increasing PHYB abundance enlarged the PB size (Fig. 1d,e). The majority of PHYB-CFP in PBs was immobile in *gPBC-25* (Fig. 1g). This attribute of PHYB condensates was similar to the biomolecular condensates formed by PHYB alone in mammalian cells^37^, suggesting that PHYB molecules were tightly bound in PBs. Surprisingly, however, further increasing the PHYB level in *PBC* dramatically reduces the fluorescence recovery kinetics and the mobile fraction of PHYB-CFP in PBs, indicative of a gel-like state (Fig. 1g). Because even in *gPBC-25* PHYB-CFP was expressed at a level 7-fold higher than that of endogenous PHYB (Fig. 1c), these results may suggest that changes in PHYB abundance under the physiological PHYB concentration may change both PB size and dynamics. Another unexpected result is that increasing PHYB abundance elicited a biphasic PIF5 response, implying that the nucleoplasmic PHYB increased in *gPBC-25* to promote PIF5 degradation. Theoretically, if PBs were formed via LLPS of PHYB, increasing PHYB in a LLPS system should only enhance the amount of PHYB in PBs but not in the nucleoplasm. One possibility is that at the physiological PHYB concentration PBs can form below the critical concentration for PHYB LLPS and therefore may require a complex mechanism beyond LLPS.

This study reveals a novel PB-mediated light signaling mechanism, in which PB formation phase-separates two opposing PHYB signaling actions of PIF5 stabilization and degradation to titrate environmental responses (Fig. 6). This PB-enabled signaling mechanism provides the framework for regulating a plethora of PHYB-interacting signaling molecules in diverse plant environmental responses. We propose that this PB function represents a general function of biomolecular condensates to allow distinct variations of a cellular process or signaling pathway to coexist and interact to generate dynamically adjustable integrated outputs within a single subcellular space.

## METHODS

### Plant materials, growth conditions, and hypocotyl measurements

The *Arabidopsis* Columbia ecotype (Col-0) was used throughout this study. The *phyB-9*^77^, *pifq* (Col-0)^78^ and *pif5-3* (SALK_087012)^79^ mutants, as well as the *PBC* line^32^, were previously described. The *gPBC-25*, *gPBC-29*, *mCherry-PIF5/pif5-3* (#*8* and *#9*) and *mCherry-PIF5/PBC* (*#1* and #*9*) transgenic lines were generated in this study. *Arabidopsis* seeds were surface-sterilized and plated on half-strength Murashige and Skoog (½ MS) medium containing Gamborg’s vitamins (MSP06, Caisson Laboratories, Smithfield, UT), 0.5 mM MES pH 5.7, and 0.8% agar (w/v). Seeds were stratified in the dark at 4°C for five days before treatment with specific light conditions and temperatures in an LED chamber (Percival Scientific, Perry, IA). Seedlings grown in the dark were exposed to 10 μmol m^-2^ s^-1^ FR light for 3 h after stratification to induce germination. Fluence rates of light were measured using an Apogee PS-200 spectroradiometer (Apogee Instruments, Logan, UT) and SpectraWiz^®^ spectroscopy software (StellarNet, Tampa, FL). Images of representative seedlings were taken using a Leica M165 FC stereo microscope (Leica Microsystems, Deerfield, IL) and processed using Adobe Photoshop CC (Adobe, San Jose, CA). For hypocotyl measurements, seedlings were scanned using an Epson Perfection V700 photo scanner (Epson America, Los Alamitos, CA), and hypocotyl length was measured using NIH ImageJ software (http://rsb.info.nih.gov/nih-image/).

### Plasmid construction and generation of transgenic lines

To generate the *gPBC* lines, genomic *PHYB* DNA with *CFP* inserted in-frame before the stop codon was cloned into *pJHA212G*^80^. The resulting construct was transformed into *phyB-9*. To generate *mCherry-PIF5/pif5-3* lines, a 2.3-kb genomic DNA fragment encompassing the *PIF5* promoter region, 5’ UTR, and the first two codons was amplified using PCR and subcloned with 4× *Myc*, *mCherry-2-L* and the *PIF5* coding sequence into *pJHA212G*^80^ containing the *rbcS* terminator using Gibson Assembly (New England Biolabs, Ipswich, MA). The resulting construct was transformed into *pif5-3* using Agrobacterium-mediated transformation. To generate *mCherry-PIF5/PBC* lines, a 1.2-kb *UBQ10* promoter was amplified using PCR and subcloned with 3× *HA*, *mCherry-2-L* and the *PIF5* coding sequence into *pJHA212B*^80^ containing an *rbcS* terminator using Gibson Assembly. The resulting construct was transformed into *PBC* Agrobacterium-mediated transformation. The primers used for plasmid construction in this study are listed in Supplementary Table 1.

### Protein extraction and immunoblotting

Total protein was extracted from *Arabidopsis* seedlings grown under the indicated conditions. To measure the degradation kinetics of PIF5 and mCherry-PIF5, seedlings were vacuum infiltrated for 15 min in PBS containing 200 μM cycloheximide and then collected at the indicated time points. At least three independent degradation kinetics experiments were performed to calculate the average relative PIF5 or mCherry-PIF5 levels at each time point normalized to the PIF5 value at time zero. Plant tissues were ground in liquid nitrogen and resuspended in extraction buffer containing 100 mM Tris-HCl pH 7.5, 100 mM NaCl, 5% SDS, 5 mM EDTA pH 8.0, 20 mM dithiothreitol, 20% glycerol, 142 mM β-mercaptoethanol, 2 mM phenylmethylsulfonyl fluoride, 1× cOmplete™ EDTA-free Protease Inhibitor Cocktail (Sigma-Aldrich, St. Louis, MO), 80 μM MG132, 80 μM MG115, 1% phosphatase inhibitor cocktail 3 (Sigma-Aldrich, St. Louis, MO), 10 mM N-ethylmaleimide (NEM), 2 mM sodium orthovanadate (Na_3_OV_4_), 25 mM β-glycerophosphate disodium salt hydrate, 10 mM NaF, and 0.01% bromophenol blue. Protein extracts were boiled for 10 min and then centrifuged at 16000L×L*g* for 10 min at room temperature. Protein extracts were separated via 12% SDS-PAGE, transferred to a PVDF membrane, probed with the indicated primary antibodies, and then incubated with HRP-conjugated secondary antibodies. Rabbit anti-PIF5 polyclonal antibody (AS12 2112, Agrisera, Vännäs, Sweden) was used at a 1:1000 dilution. Mouse anti-PHYB monoclonal antibody (a gift from Dr. Akira Nagatani) was used at a 1:2000 dilution. Goat anti-mouse (1706516, Bio-Rad Laboratories, Hercules, CA) and goat anti-rabbit (1706515, Bio-Rad Laboratories, Hercules, CA) secondary antibodies were used at a 1:5000 dilution. Signals were detected via SuperSignal West Dura Extended Duration Chemiluminescent Substrate (Thermo Fisher Scientific, Waltham, MA).

### Immunofluorescence staining

Whole-mount immunofluorescence staining was performed as described previously with the following modifications^73,81^. Seedlings were fixed in 4% (v/v) paraformaldehyde (15710, Electron Microscopy Sciences, Hatfield, PA), dehydrated, and mounted on slides^73,81^. All subsequent steps were performed in a 55-µL SecureSeal chamber (621505, Grace Bio-Labs, Bend, OR). Myc-tagged mCherry-PIF5 was detected using rabbit anti-Myc polyclonal antibodies (2272, Cell Signaling Technology, Danvers, MA) as the primary antibody at 1:100 dilution and donkey anti-rabbit AlexaFluor 555 antibodies (A31572, Thermo Fisher Scientific, Waltham, MA) as the secondary antibody at 1:1000 dilution. PHYB was detected using mouse monoclonal anti-PHYB antibodies (a gift from Akira Nagatani, 1:100 dilution) as the primary antibody and donkey anti-mouse AlexaFluor 488 antibodies (A21202, Thermo Fisher Scientific, Waltham, MA, 1:1000 dilution) as the secondary antibody. Nuclei were counterstained with 3.6 µM 4′,6-diamidino-2-phenylindole (DAPI). Samples were mounted using ProLong Gold Antifade Mountant (Thermo Fisher Scientific, Waltham, MA) and left to cure overnight in the dark before confocal analysis.

### Seedling preparation for PB analysis using confocal microscopy

Seedlings were fixed following a previously described protocol with slight modifications^82^. Seedlings were fixed under vacuum with 1% (v/v) paraformaldehyde in PBS for 10 min. After quenching with 50LmM NH_4_Cl, the fixed seedlings were permeabilized with 0.2% Triton X-100 in PBS, and nuclei were stained with 3.6 µM DAPI in PBS for 10 min. Seedlings were washed with PBS before being mounted on a slide using Prolong Diamond Antifade Mountant (Thermo Fisher Scientific, Waltham, MA). The slides were left to cure overnight in the dark before being sealed with nail polish and stored at 4°C. For live-cell imaging, seedlings were mounted using PBS. The slides were transported to the microscope in an aluminum foil-wrapped petri dish, and nuclei were imaged within 5 min after mounting.

### Fluorescence microscopy and imaging analysis

For colocalization of mCherry-PIF5 and PHYB in *mCherry-PIF5/pif5-3* using immunofluorescence staining, three-dimensional (3D) image stacks of individual nuclei of cotyledon pavement epidermal cells were imaged using a Zeiss LSM800 confocal microscope equipped with a ×100/1.4 Plan-Apochromat oil-immersion objective (Carl Zeiss AG, Jena, Germany). Alexa 488 was monitored using 488 nm excitation and 490-561 nm bandpass emission.. Alexa 555 was monitored using 561 nm excitation and 560-630 nm bandpass emission. DAPI was monitored using 405 nm excitation and 410-470 nm bandpass emission. For colocalization mCherry-PIF5/PBC lines, 3D image stacks of individual nuclei of cotyledon pavement epidermal cells were imaged using a Zeiss LSM800 confocal microscope equipped with a ×100/1.4 Plan-Apochromat oil-immersion objective (Carl Zeiss AG, Jena, Germany). PHYB-CFP was monitored using 405 nm excitation and 410-470 nm bandpass emission. mCherry-PIF5 was monitored using 561 nm excitation and 560-630 nm bandpass emission. DAPI was monitored using 405 nm excitation and 410-470 nm bandpass emission. Deconvolution (nearest neighbor) was performed and the maximum projection of image stacks (Z-stack interval 0.7 μm) was generated using Zeiss ZEN 2.3 software (Carl Zeiss AG, Jena, Germany). Representative images were exported as TIFF files and processed using Adobe Photoshop CC (Adobe, San Jose, CA). To measure the PB diameter of fixed *mCherry-PIF5/PBC* seedlings, the boundary of the PB peak was defined as the point where the fluorescence intensity was half of the peak value.

For PB analysis in *gPBC-25*, *gPBC-29*, *PBC*, and dark grown *mCherry-PIF5/PBC*, 3D image stacks of individual nuclei from cotyledon epidermal or upper hypocotyl epidermal cells were imaged using a Zeiss Axio Observer Z1 inverted microscope equipped with a Plan-Apochromat 100x/1.4 oil-immersion objective and an Axiocam mono camera (Carl Zeiss, Jena, Germany). Fluorescence was detected using a broad spectrum X-Cite 120LED Boost high-power LED illumination system (Excelitas Technologies, Walham, MA) and the following Zeiss filter sets: DAPI, excitation 365 nm, emission 445/50 nm/nm (Zeiss Filter Set 49); CFP, excitation 436/25 nm/nm, emission 480/40 nm/nm (Filter set 47 HE); mCherry, excitation 550/25 nm/nm, emission 605/70 nm/nm (Zeiss Filter set 43 HE), brightfield was used for the DIC image. Image stacks with a Z-step size of 0.24 µm were subjected to iterative classic maximum likelihood estimation deconvolution using Huygens Essential (Scientific Volume Imaging, Hilversum, Netherlands). The volume of PBs was quantified using Huygens Object Analyzer (Scientific Volume Imaging, Hilversum, Netherlands). The PB partitioning of PHYB-CFP was measured using Huygens Object Analyzer and 3D image stacks of PHYB-CFP and the DAPI-stained nucleus. Huygens Object Analyzer renders a 3D model of the PHYB-CFP signals in PBs and the entire nucleus that was defined by the DAPI signal. The PB-partitioning of PHYB-CFP was calculated as the percentage of PHYB-CFP in PBs versus the entire nucleus.

For FRAP experiments, seedlings were mounted in PBS and placed onto a Zeiss LSM800 confocal microscope with a C-Apochromat 63×/1.2 W autocorr M27 objective (Carl Zeiss AG, Jena, Germany). For the *gPBC-2*5 and *PBC* experiments, PHYB-CFP was monitored using 405 nm excitation and 490-561 nm bandpass detector settings. Images were acquired at 2.7× magnification, the image format was 512×512 pixels, and the pinhole was set at 97 μm. One image was taken pre-bleach, followed by 6 iterations of photobleaching using the 405 nm laser at 85% power. Images after bleaching were collected using low laser intensities and were taken at 4 s intervals for 18 cycles. For the *mCherry-PIF5/PBC* experiments, PHYB-CFP was monitored using 405 nm excitation and 490-561 nm bandpass detector settings, and mCherry-PIF5 was excited with a 561 nm laser and observed using a 560-630 nm bandpass emission. Images were acquired at 2.7× zoom, the image format was 512×512 pixels, and the pinhole was 121 μm. One image was taken pre-bleach, followed by 8 iterations of photobleaching using the 405 nm laser at 50-60% power for PHYB-CFP and the 561 nm laser for mCherry-PIF5 at 85% power. Images after bleaching were collected using low laser intensities and were taken at 6 s intervals for 20 cycles. FRAP curve analyses were performed by setting the time of bleach to zero. Normalization was performed by taking the sum of intensity of the ROI, subtracting the background, and setting the initial pre-bleach intensity to 1. Data were fitted to a one-phase exponential curve, and MF and t_1/2_ were calculated using Prism 10 (GraphPad Software, Boston, MA).

### RNA extraction and quantitative real-time PCR

Total RNA from seedlings was isolated using a Quick-RNA MiniPrep kit with on-column DNase I treatment (Zymo Research, Irvine, CA). cDNA was synthesized using Oligo(dT) primers and 1 μg total RNA via a Superscript II First-Strand cDNA Synthesis Kit (Thermo Fisher Scientific, Waltham, MA). Quantitative real-time PCR (qRT-PCR) was performed with iQ SYBR Green Supermix (Bio-Rad Laboratories, Hercules, CA) using a LightCycler 96 System (Roche, Basel, Switzerland). The transcript level of each gene was normalized to that of *PP2A* (At1G13320). The statistical analyses were performed using one-way analysis of variance (ANOVA) with post hoc Tukey’s HSD using Prism 10 (GraphPad Software, Boston, MA). The primers for qRT-PCR analysis used in this study are listed in Supplementary Table 2.

## DATA AVAILABILITY

*Arabidopsis* mutants and transgenic lines as well as plasmids generated during the current study are available from the corresponding author upon reasonable request. Source data are provided with this paper.

## ACKNOWLEDGEMENTS

We thank Akira Nagatani (Kyoto University) for providing anti-PHYB antibodies. We thank Elise Pasoreck for valuable suggestions and comments on the manuscript. We are grateful to Xuemei Chen for her support of D.F. in a collaborative project on thermomorphogenesis. This work was supported by National Institute of General Medical Sciences grant R01GM087388 to M.C.

## AUTHOR CONTRIBUTIONS

R.J.K., D.F., J.H., K.K., J.D. and M.C. conceived of the original research plan; M.C. supervised the experiments; R.J.K., D.F., J.H., K.K., J.D. and M.C. performed the experiments; R.J.K., D.F., J.H., K.K., J.D. and M.C. analyzed the data; R.J.K., D.F., J.H., and M.C. wrote the article with contributions from all authors.

## SUPPLEMENTARY INFORMATION

**Supplementary Table 1. List of primers used for plasmid construction.**

**Supplementary Table 2. List of primers used for qRT-PCR experiments.**

